# PIK3IP1/TrIP restricts activation of T cells through inhibition of PI3K/Akt

**DOI:** 10.1101/214981

**Authors:** Uzodinma Uche, Andrea L. Szymczak-Workman, Stephanie Grebinoski, Ann Piccirillo, Louise M. D’Cruz, Lawrence P. Kane

**Affiliations:** Department of Immunology; Graduate Program in Microbiology and Immunology University of Pittsburgh School of Medicine Pittsburgh, PA 15261

## Abstract

Phosphatidylinositol-3 kinases (PI3Ks) modulate numerous cellular functions, including growth, proliferation and survival. Dysregulation of the PI3K pathway can lead to autoimmune disease and cancer. PIK3IP1 (or Transmembrane Inhibitor of PI3K – TrIP) is a novel transmembrane regulator of PI3K. TrIP contains an extracellular kringle domain and an intracellular “p85-like” domain with homology to the inter-SH2 domain of the regulatory subunit of PI3K. Although TrIP has been shown to bind to the p110 catalytic subunit of PI3K in fibroblasts, the mechanism by which TrIP functions is poorly understood. We show that both the kringle and “p85-like” domains are necessary for TrIP inhibition of PI3K. We also demonstrate that TrIP protein is down-modulated from the surface of T cells to allow T cell activation. In addition, we present evidence that the kringle domain may modulate TrIP function by binding an as-yet-unidentified ligand. Using an inducible knockout mouse model that we developed, we show that TrIP-deficient T cells exhibit more robust T cell activation, show a preference for Th1 polarization and can mediate clearance of *Listeria monocytogenes* infection faster than WT mice. Thus, TrIP is an important negative regulator of T cell activation and may represent a novel target for immune modulation therapies.

## INTRODUCTION

Phosphatidylinositide-3-Kinases (PI3K) are a family of lipid kinases that play important intracellular signaling roles in cellular processes such as proliferation, motility, growth, intracellular trafficking, differentiation and survival (Cantley, 2002;; Fruman, 2007;; Han et al., 2012). There are three main classes of PI3K. Class I PI3Ks, which are prevalent in immune cells, are composed of two subunits: a regulatory subunit (p85) and a catalytic subunit (p110) (Engelman, 2009;; Fresno Vara et al., 2004;; Fruman et al., 1998). During T cell receptor activation, PI3K is recruited to the plasma membrane via the SH2 domain of the p85 subunit. The associated p110 subunit is then activated to phosphorylate phosphatidylinositol 4,5-bisphosphate (PIP_2_) and produce phosphatidylinositol (3,4,5)-trisphosphate (PIP3). PIP_3_ interacts with the pleckstrin homology domain of Akt, causing a conformational change that allows PDK1 (kinase 3-phosphoinositide-dependent protein kinase-1) to partially activate Akt by phosphorylating threonine 308 (T308). Full activation of Akt is achieved by mTORC2-mediated phosphorylation at serine 473 (S473) and facilitates such processes as cell growth, cell cycle progression and cell survival. It is therefore not surprising that Akt amplification due to dysregulation of PI3K has been implicated in many cancers. In fact, several studies are presently focused on the development of PI3K pathway inhibitors as a means of combating various forms of cancer (Engelman, 2009).

Several negative regulators of PI3K have been identified (Agoulnik et al., 2011;; Antignano et al., 2010;; Carracedo and Pandolfi, 2008;; Dillon and Miller, 2014). PTEN (phosphatase and tensin homologue deleted on chromosome 10) and SHIP-1 (SH2 containing inositol 5’-phosphatase) are phosphatases that dephosphorylate PIP_3_ to PIP_2_, thereby inhibiting downstream signaling in the PI3K pathway. INPP4B (inositol polyphosphate 4-phosphatase type II) has been shown to dephosphorylate PIP_2_, thereby playing a role in the negative regulation of the PI3K pathway. Several studies have shown that loss of function mutations or deletions of these phosphatases can lead to dysregulated PI3K activity.

While the above phosphatases act downstream of PI3K, PIK3IP1 (PI3K-interacting protein- 1, which we will refer to as TrIP (transmembrane inhibitor of PI3K) for simplicity) is a recently identified inhibitor that acts upstream of the aforementioned phosphatases (DeFrances et al., 2012;; Zhu et al., 2007). TrIP is a transmembrane protein composed of two main domains, an extracellular kringle domain and an intracellular tail that includes a motif similar to the p110-binding inter-SH2 domain found in the p85 subunit of PI3K. Overexpression of TrIP in mouse hepatocytes leads to a reduction in PI3K signaling and suppression of hepatocyte carcinoma development (He et al., 2008).

Furthermore, recent work in cancer genetics highlights the transcriptional down-regulation of TrIP as a contributing factor to dysregulated PI3K signaling in tumorigenesis (Wong et al., 2014). Although it has been shown that TrIP inhibits PI3K by binding the p110 subunit via the p85-like domain, the role of the kringle domain remains to be determined. Given the ability of kringle domains in other proteins to bind to various ligands, it is possible that the TrIP kringle domain may bind one or more ligands for modulation of TrIP activity (Christen et al., 2010;; Mikels et al., 2009;; Patthy et al., 1984). Because TrIP is highly expressed in immune cells, particularly mast cells and T cells (DeFrances et al., 2012), we wanted to investigate how the structure of TrIP enables regulation of PI3K in the context of the activated T cell.

In this study, we investigated the importance of both the kringle and p85-like domains to TrIP function in activated T cells. We also examined how cell fate decisions and immune response are regulated by TrIP. Here we show that both the extracellular kringle domain and the intracellular p85- like domain are necessary for inhibition of PI3K by TrIP. Intriguingly, we also show that cell-surface levels of TrIP are decreased upon T cell activation, which correlates with the upregulation of PI3K pathway signaling. Using a T cell conditional knockout mouse model, we show that the loss of TrIP in T cells leads to an increase in T cell activation, which translates to stronger Th1 inflammatory potential and more rapid clearance of *L. monocytogenes* infection.

## MATERIALS AND METHODS

### Cell Lines and transfections

The D10 Th2 T cell clone (D10.G4.1;; ATCC TIB-224) were maintained in RPMI media supplemented with 10% bovine growth serum (BGS), penicillin, streptomycin, glutamine and 25U/ml recombinant human IL-2. Human embryonic kidney (HEK) 293 cells were maintained in DMEM media supplemented with 10% bovine growth serum (BGS), penicillin, streptomycin and glutamine. CH27 cells (mouse lymphoma;; RRID:CVCL_7178) were maintained in RPMI media supplemented with 10% bovine growth serum (BGS), penicillin, streptomycin and glutamine. For structure-function assays, using a BIO-RAD GenePulser Xcell, D10 cells were individually electroporated with control plasmid, flag tagged wt-TrIP or flag tagged mutant TrIP constructs, along with pMaxGFP plasmid (encoding GFP from copepod *Pontellina p*.). One day after transfection, cells were evaluated by flow cytometry for GFP expression and TrIP expression (via anti-flag staining) using anti-DYKDDDDK APC clone L5 (BioLegend cat# 637308). For p110δ interaction assays, flag-tagged TrIP variants and HA-tagged p110δ were transfected into HEK293 cells using TransIT-LT1 (Mirus) according to the manufacturers protocol. Cells were evaluated by western blot for expression using Roche anti-HA (clone 12CA5;; Cat# 11 583 816 001) and BioLegend Direct-Blot HRP anti-DYKDDDDK (Clone L5;; cat# 637311).

### T cell/APC co-cultures

CH27 cells were pulsed with 100 µg/ml chicken conalbumin (Sigma C7786) one day before mixing with D10 cells. D10 cells were mixed in a 1:1 ratio with either pulsed unpulsed CH27 cells for 15, 30 and 60 mins. Reactions were quenched by placing cells on ice, adding 1 ml of PBS and centrifugation to settle cells and decant activation media. Cells were then stained with anti-DYKDDDDK APC clone L5 (BioLegend cat# 637308), anti-mouse CD19 violetFluor 450 (clone 1D3;; Tonbo Biosciences 75-0193-U025) and anti-mouse CD4 Brilliant Violet 510 (clone GK1.5;; BioLegend #100449). Cells were washed three times in PBS and then fixed and permeabilized with eBioscience Foxp3/Transcription factor staining buffer (cat # 00-5523-00), then stained with anti-pS6 (S235/236) Alexa 647 (clone D57.2.2E;; Cell Signaling #5316S).

### Mice

Mice with a floxed *Pik3ip1* gene (Pik3ip1^fl/fl^) were generated by inGenious Targeting Laboratory Inc., using C57BL/6 ES cells. CD4-Cre mice on a C57BL/6 background were originally purchased from Taconic, then maintained by breeding to C57BL/6J mice (Jackson), and were used as controls. All mouse strains were on a C57BL/6 background and maintained in facilities of the University of Pittsburgh Division of Laboratory Animal Resources. Mice were age-matched within experiments, with approximately equal numbers of male and female animals. All mouse studies were performed in accordance with University of Pittsburgh Institutional Animal Care and Use Committee procedures.

### T cell purification and differentiation

CD4 T cells from spleens and lymph nodes of naïve mice were purified by magnetic separation using the Miltenyi Biotec naïve CD4 T cell isolation kit (Cat# 130-104-453). The purity of the final cell population was >90%. T cells were activated with plate-bound anti-CD3 (clone 145-2C11;; Bio X Cell InVivoMab BE0001-1) in complete (RPMI medium supplemented with 10% bovine growth serum, 2 mM L-glutamine, 100U/ml penicillin, 100 µg/ml streptomycin, 50 µM 2-mercaptoethanol, HEPES and sodium pyruvate). For Th1 differentiation, cells were cultured in the presence of recombinant mouse IL-12 (10 ng/ml), anti-IL-4 (10 µg/ml) and recombinant human IL-2 (50 U/ml). For Th17 differentiation, cells were cultured in the presence of recombinant human TGF-B (2.5 ng/ml), recombinant mouse IL-6 (20 ng/ml), and anti-IFNγ (10 µg/ml), anti-IL-4 (10 µg/ml), anti-IL-2 (20 µg/ml) neutralizing antibodies. For iTreg differentiation, cells were cultured in the presence of anti-IL-4 (10 µg/ml) and anti-IFNγ (10 µg/ml) neutralizing antibodies as well as recombinant human TGFB2 (10 ng/ml) and recombinant human IL-2 (50 U/ml). Cells were cultured for 3 days under Th differentiation conditions and then stimulated for 4h with PMA (50ng/ml) and ionomycin (1.33µM) in the presence of golgi plug (BD biosciences cat#51-2301KZ). Cells were then harvested and analyzed by flow cytometry.

T helper cell differentiation reagents: Mouse IL-12 (Miltenyi Biotec Cat# 130-096-707). Mouse IL-6 (Miltenyi Biotec Cat# 130-094-065). TGF-β human recombinant (Sigma SRP3170-5UG). Anti-mouse IL-4 clone 11B11(Bio X Cell #BE0045). Anti-mouse IL-2 clone S4B6-1 (Bio X Cell #BE0043). Anti-mouse IFN-γ clone XMG1.2 (Bio X Cell #BE0055).

The following antibodies and dyes were used for flow cytometry: Anti-mouse CD4 Brilliant Violet 510 clone GK1.5 (BioLegend #100449). Ghost Red 780 viability dye (TONBO biosciences #13-0865-T100). V450-rat anti-mouse Foxp3 Clone MF23 (BD Horizon #561293). mouse anti-mouse RORγt PE (clone Q31-378; BD Biosciences 562607). Anti-mouse IL-17A PerCP-Cy5.5 (clone eBio17B7;; eBioscience 45-7177-80). Anti-mouse Foxp3 APC (clone FJK-16s;; eBioscience 17-5773-82). Anti-mouse T-bet eFluor 660 (clone eBio4B10;; eBioscience 50-5825-82). Anti-mouse IFN-γ PE-Cy7 (clone XMG1.2;; Tonbo Biosciences 60-7311-U025). Anti-mouse CD25 FITC (clone PC61.5; Tonbo Biosciences 35-0251-U100).

### T cell activation

Splenocytes and lymphocytes obtained from conditional KO and WT mice were stimulated in complete RPMI with 3 µg/ml biotinylated anti-CD3 and anti-CD28 in the presence of 15 µg/ml streptavidin (anti-mouse CD3e biotin, clone 145-2C11, Tonbo Biosciences #30-0031-U500;; hamster anti-mouse CD28 biotin, clone 37.51, BD Biosciences #553296;; streptavidin, cat#189730, MilliporeSigma). Cells were activated for 15 mins, 30 mins, 60 mins or 4 hours. Activation was stopped by placing cells on ice, adding 1 ml of cold PBS and centrifugation, to settle cells, followed by aspiration of media.

The following antibodies and dyes were used for flow cytometry: Anti-mouse CD4 Brilliant Violet 510 (clone GK1.5;; BioLegend #100449). Anti-mouse CD8a v450 (clone 53-6.7;; Tonbo Biosciences 75-0081-U100). Anti-pS6 (S235/236) Alexa 647 (clone D57.2.2E;; Cell Signaling #5316S). Hamster anti-mouse CD69 FITC (clone H1.2F3;; BD Biosciences #557392). Ghost Red 780 viability dye (Tonbo Biosciences #13-0865-T100).

### qPCR

RNA was extracted using the Qiagen RNeasy Mini Kit (cat# 74106) and reverse-transcribed to generate cDNA with the Applied Biosystems High Capacity cDNA Reverse Transcription Kit (cat# 4368813). Quantitative real-time polymerase chain reaction assays were performed with Power SYBR Green PCR Master Mix (Applied Biosystems cat# 4367659) on a Step-One Plus Real Time PCR system. The abundance of TrIP mRNA was normalized to that of mGAPDH as calculated with the 2^−ΔΔCT^ method. The primers used are as follows: Forward: 5’-ATGCTGTTGGCTTTGGGTACAC-3’ Reverse: 5’-CGGCAGTAGTTGTGGTTGC-3’

### Flow cytometry

Before staining, cells were washed in staining buffer (1% bovine growth serum supplemented PBS). For extracellular staining, cells were stained at 4˚C with antibodies resuspended in staining buffer. For intracellular staining, cells were washed three times to remove excess extracellular staining and then fixed and permeablized with either the eBioscience Foxp3/Transcription factor staining buffer set (cat # 00-5523-00) or the BD Biosciences cytofix/perm kit (cat# 554714) for transcription factor and cytosol analysis, respectively. Fixed and permeablized cells were then stained at 4˚C with antibodies resuspended in 1X permeabilization buffer from the respective kits.

### Western blotting analysis and immunoprecipitations

Cells were lysed in ice-cold NP-40 lysis buffer (1% NP-40, 1mM EDTA, 20mM tris-HCL (pH 7.4), 150mM NaCl) for both protein analysis and immunoprecipitation. Immunoprecipitation was performed by mixing lysate with 20µl of M2 anti-flag agarose beads for 3 hrs at 4 deg. C. Proteins were eluted from beads by mixing Lysate with 1X SDS (containing 5% B-mercaptoethanol) and boiling at 95 deg. C for 10 mins. For western blot analysis, proteins were resolved by 10% SDS-polyacrylamide gel electrophoresis and were transferred onto polyvinylidene difluoride membranes which were then blocked in 4% BSA. The membranes were then incubated with the primary antibodies overnight. This was followed by incubating the membrane with HRP-conjugated secondary antibodies for 2 hours before detection with the SuperSignal West Pico ECL substrate (Thermo Fisher Scientific) and imaging on a Protein Simple FluorChem M.

*Western blot and CoIP reagents:* Anti-HA (12CA5) (Roche Cat# 11-583-816-001). Anti-Flag M2 Affinity Gel (Sigma Cat# A2220). Direct-Blot HRP anti-DYKDDDDK (Clone L5;; BioLegend cat#637311).

### Listeria infection

Control (CD4-Cre alone) and Pik3ip1^fl/fl^ x CD4-Cre mice were infected intravenously with 15,000 CFU LM-GP33 resuspended in 200µl of PBS. Bacterial titers were quantified by lysing of whole livers in PBS and plating of a 10 fold dilution of bacteria on brain-heart infusion agar plates overnight. Immune response to LM-GP33 infection was quantified by extracting splenocytes and staining with a GP33 specific tetramer in complete RPMI at room temperature for 1 hour. After Tetramer staining, cells were stained with the following antibodies and dyes and analyzed by flow cytometry: Ghost Red 780 viability dye (Tonbo Biosciences #13-0865-T100), anti-mouse CD4 Brilliant Violet 510 (clone GK1.5; BioLegend #100449), anti-mouse CD8a PE (clone 53-6.7; Tonbo Bioscience 50-0081-U100), anti-mouse CD62L PerCP-Cy5.5 (clone MEL-14; eBioscience 45-0621-80), Anti-mouse KLRG1 FITC (clone 2F1; Tonbo Biosciences 35-5893-U100), anti-mouse CD127 PE-Cy7 (clone A7R34; Tonbo Biosciences 60-1271-U025), and anti-mouse CD44 violetFluor 450 (clone IM7; Tonbo Biosciences 75-0441-U025). All Ab staining was performed at 4˚C in PBS containing 1% BGS and 0.1% sodium azide. GP33 tetramers were provided as APC fluorophore-conjugated tetramers by the NIH tetramer core facility and used for identification of gp33-specific CD8^+^ and CD4^+^ T cells.

## RESULTS

### TrIP inhibits PI3K/mTOR pathway signaling

We have previously shown that ectopic expression of TrIP in T cells can inhibit the phosphorylation of Akt and thus its activation (DeFrances et al., 2012). A more sensitive readout of PI3K/Akt/mTOR activity is analysis of ribosomal protein S6 phosphorylation (pS6) by flow cytometry. In order to study the structural requirements of TrIP for modulation of T cell function, we evaluated pS6 in D10 T cells, a cell line with apparently normal PI3K signaling (Kane et al., 2004), in the context of ectopically expressed WT or mutant TrIP. The domain structure of TrIP is illustrated in **Fig. 1A**. In the absence of a suitable antibody for detecting TrIP by flow cytometry, we transfected D10 T cells with control plasmid or Flag-tagged TrIP (WT TrIP) and monitored TrIP expression using α-Flag antibody (**Fig. 1B**). Co-transfection of GFP was also used to determine transfection efficiency. One day after transfection, cells were stimulated with α-CD3/CD28 and analyzed at various time points (**Fig. 1C-D**). At all time points evaluated, cells with ectopic TrIP expression displayed lower pS6 compared with empty vector-transfected cells. We also noted that, in the absence of stimulation, ectopic expression of TrIP resulted in lower basal pS6. These results support previous data from our lab and others suggesting that TrIP inhibits Akt activation in T cells (DeFrances et al., 2012; Menhart et al., 1995; Mikels et al., 2009; Patthy et al., 1984).

**Figure 1.**
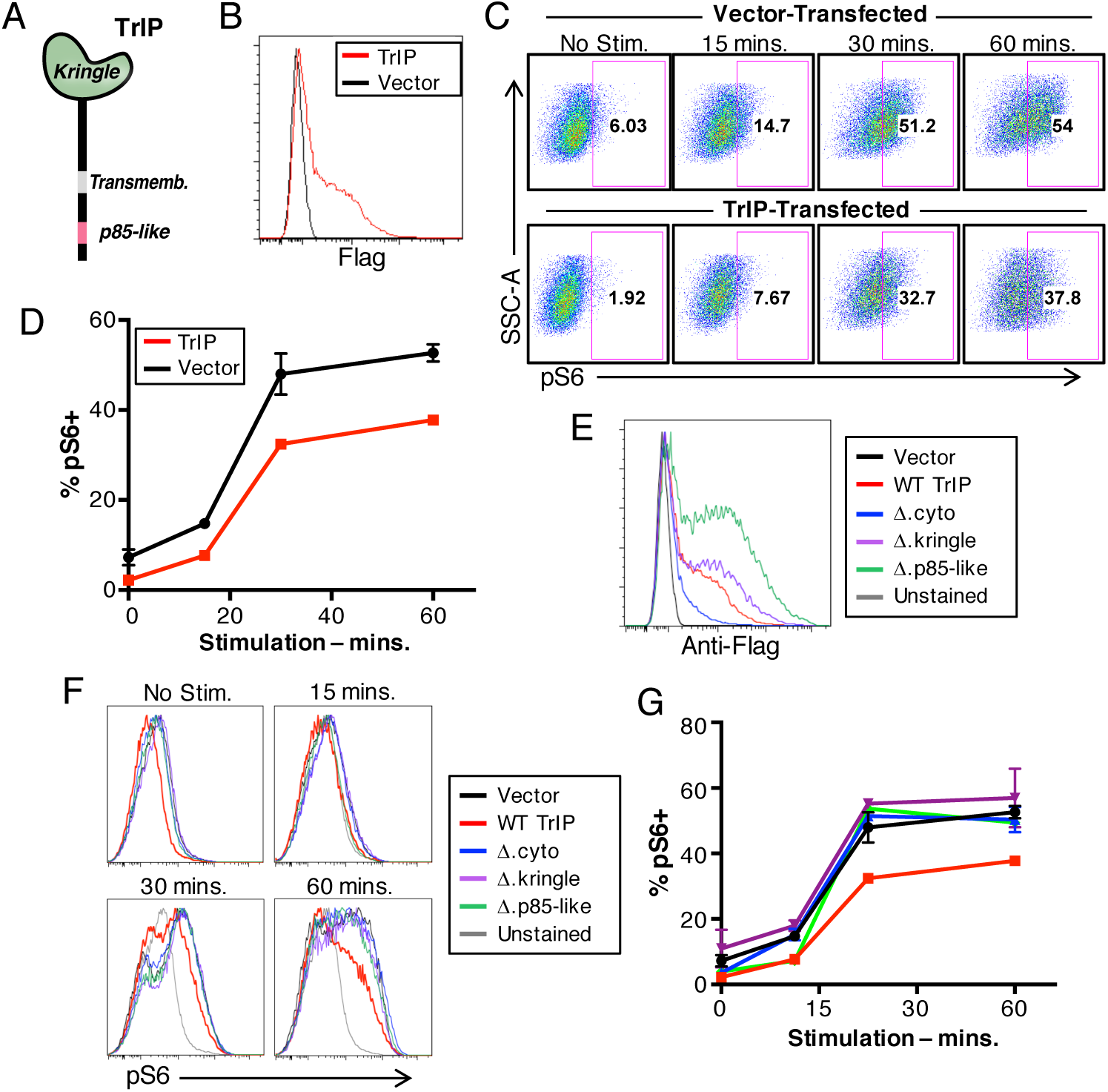
The kringle and cytoplasmic domains are required for TrIP activity. (**A**) Domain structure of TrIP, indicating the kringle, transmembrane and p85-like domains. (**B**) Expression of Flag-tagged WT TrIP on transiently transfected D10 T cells. (**C**) Flow cytometric analysis of pS6 activation in control (vector) or WT TrIP-transfected D10 T cells, stimulated with anti-CD3/CD28 for the indicated times. (**D**) Quantitation of pS6 activation as shown in panel C. (**E**) Expression of WT and mutant Flag-tagged TrIP constructs on transfected D10 cells. (**F**) Representative histogram of pS6 staining in D10 cells transiently transfected with Flag-tagged TrIP constructs and activated with anti-CD3/CD28. (**G**) Quantitation of pS6 activation obtained from flow cytometric analysis of activated D10 cells transfected with Flag-tagged TrIP constructs, as shown in panel F. Data in each panel are representative of at least three experiments.

### The kringle and cytoplasmic domains are required for TrIP activity

The importance of the kringle domain as a ligand-binding domain in other proteins (Menhart et al., 1995; Mikels et al., 2009; Patthy et al., 1984), and the degree of homology between the p85-like domain and the inter-SH2 domain of the p85 regulatory subunit of PI3K, suggests that these two domains may play important roles in the inhibition of PI3K by TrIP. To address this, we designed Flag-tagged TrIP variants lacking either the extracellular kringle domain (Δkringle-TrIP), the entire cytoplasmic region (Δcyto-TrIP) or only the p85-like domain (Δp85-like-TrIP). These constructs were transfected, along with a GFP transfection control, into D10 T cells (**Fig. 1E**) which were then stimulated with α-CD3/CD28 and analyzed by flow cytometry for S6 phosphorylation (**Fig. 1F-G**).

While WT TrIP led to the attenuation of pS6, deletion of either the p85-like domain alone <u>or</u> the entire cytoplasmic region abolished the ability of TrIP to suppress pS6. Interestingly, expression of the kringle domain-deleted construct (which still possesses the p85-like domain) did not attenuate pS6. These results suggest that both the kringle and p85-like domains are essential for optimal TrIP inhibitory function.

### Cell-surface TrIP is downregulated upon T cell activation

Our data strongly support an inhibitory role for TrIP in T cells. Certain other negative regulators of T cell signaling (e.g. PTEN) are actively downregulated to promote T cell activation (Hawse et al., 2015; Newton and Turka, 2012), so we examined how TCR activation might affect expression of TrIP protein. In order to replicate T cell stimulation in the presence of an antigen presenting cell (APC), we used the mouse B cell lymphoma line CH27 (Haughton et al., 1986) to stimulate D10 cells. CH27 cells were either pulsed with cognate antigen (chicken conalbumin) or left unpulsed, and then mixed with D10 T cells transfected with Flag-tagged PI3KIP1. There was a notable decrease over time in Flag-TrIP expression on the surface of D10 cells (**Fig. 2A**-B). Interestingly, this decreased expression corresponded with an increase of pS6 expression (**Fig. 2C**-D). As observed when D10 T cells were stimulated with α-CD3/CD28 (**Fig. 1**), antigen-pulsed CH27 cells induced a more immediate and robust pS6 signal in T cells transfected with empty vector, compared to those transfected with WT TrIP (not shown). These results support the model that T cells acutely modulate the expression of TrIP to promote the activation of TCR-induced PI3K signaling.

**Figure 2.**
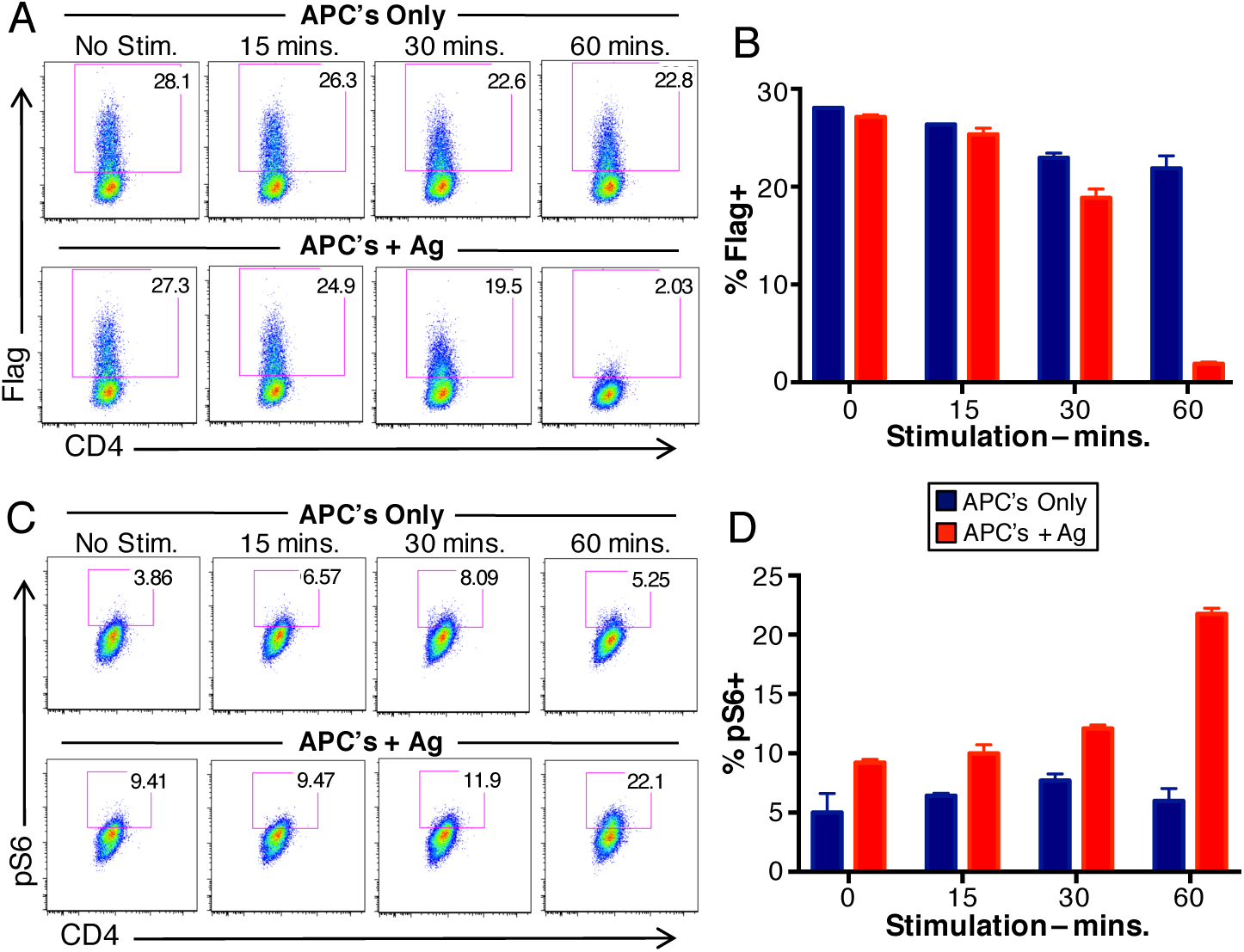
Cell-surface TrIP is downregulated during T cell activation. D10 cells transfected with WT TrIP were mixed with CH27 B cells as APCs (+/- pre-loading with conalbumin antigen) for the given time points. (**A**) Representative flow cytometry analyzing cell-surface expression of Flag-tagged WT TrIP on D10 cells stimulated with CH27 B cells alone (top row) or plus antigen (bottom row). (**B**) Quantitation of data shown in panel A. (**C**) Representative flow cytometry analyzing pS6 staining in D10 cells transfected with Flag-tagged WT TrIP and stimulated with CH27 B cells alone (top row) or plus antigen (bottom row). (**D**) Quantitation of data shown in panel C. Data are representative of three experiments.

### Structure/function analysis of TrIP cell surface expression and inhibitory activity

In order to probe the relevance of the kringle domain for TrIP expression and function, we repeated the CH27-D10 experiments with both Flag-tagged WT TrIP and ∆kringle-TrIP-transfected D10 T cells and monitored Flag-TrIP expression and S6 phosphorylation. Since ∆kringle-TrIP transfected cells showed a recovery in TCR-dependent pS6 (**Fig. 1F-G**), we suspected that the absence of the kringle domain would lead to maintenance (and not downregulation) of TrIP. As shown above, D10 cells transfected with WT TrIP and stimulated with CH27 B cells alone (without antigen) did not show significant loss of TrIP expression over time. However, when mixed with antigen-pulsed CH27 cells, WT TrIP-expressing D10 T cells showed a significant loss of TrIP expression (**Fig. 3A,** top row and **3B**). In contrast, D10 cells transfected with the Δkringle-TrIP mutant did not lose surface TrIP expression when stimulated with CH27 cells (with or without antigen) (**Fig. 3A,** bottom row and **3B**). Consistent with the results above, upon mixing with antigen-pulsed CH27 cells, ∆kringle-TrIP-expressing D10 T cells actually showed a more rapid increase in pS6 than cells transfected with WT TrIP (**Fig. 3C-D**). These results further confirm that the kringle domain is important for the inhibitory function of TrIP and suggest the possibility that TrIP is regulated by interaction with a ligand.

**Figure 3.**
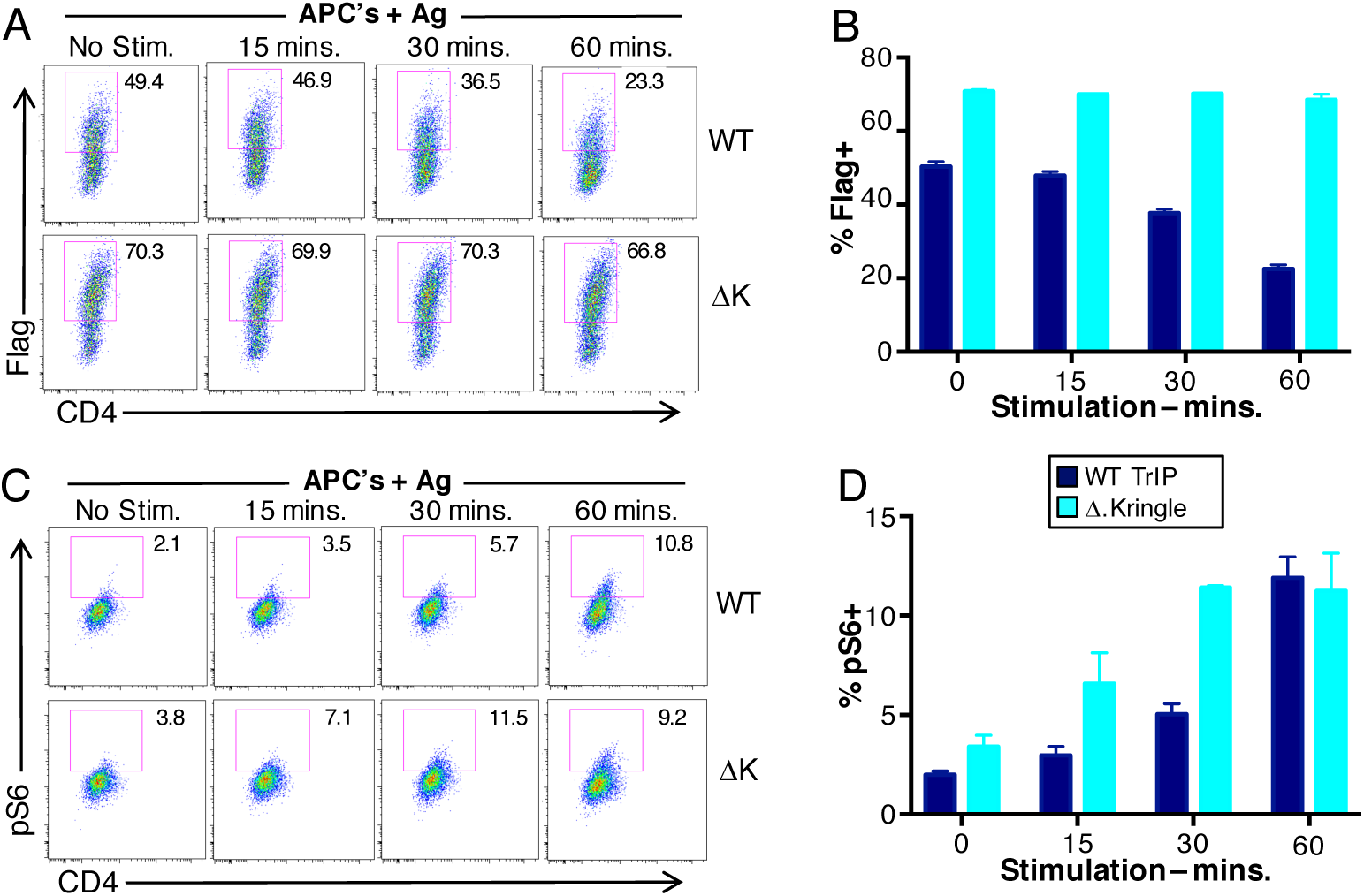
The kringle domain regulates TrIP expression and function. D10 cells transiently transfected with Flag-tagged WT or Δkringle TrIP were mixed with CH27 B cells plus antigen for the given time points. (**A**) Representative flow cytometry analyzing cell-surface expression of Flag-tagged WT (top row) or Δkringle (bottom row) TrIP on D10 cells stimulated with CH27 B cells plus antigen. (**B**) Quantitation of data shown in panel A. (**C**) Representative flow cytometry analyzing pS6 staining in the same cells as in panel A. (**D**) Quantitation of data shown in panel C. Data are representative of three experiments.

In the absence of a known ligand, we designed a hCD8-TrIP chimera to investigate possible effects of ligand engagement on TrIP function. The extracellular and transmembrane regions of human CD8 were fused with the cytoplasmic tail of TrIP, expressed in D10 T cells and detected at the cell surface with hCD8 antibody (**Fig. 4A**). Using a luciferase reporter driven by the NFAT promoter, we observed that expression of hCD8-TrIP on D10 T cells was not sufficient to inhibit TCR signaling (**Fig. 4B**). However, upon crosslinking of hCD8-TrIP with varying concentrations of α- hCD8, we observed a decrease in anti-CD3/CD28-induced pS6 (**Fig. 4C-D**). This was in contrast to the effects of a previously described CD8-CD3 ζ chimeric construct (Irving and Weiss, 1991), which upon crosslinking with anti-hCD8, modestly enhanced pS6 expression (**Fig. 4C-D**). These results suggest that binding of ligand to the kringle-domain may regulate TrIP function, possibly by aggregation of the protein and increased sequestration of PI3K from the TCR signaling machinery.

**Figure 4.**
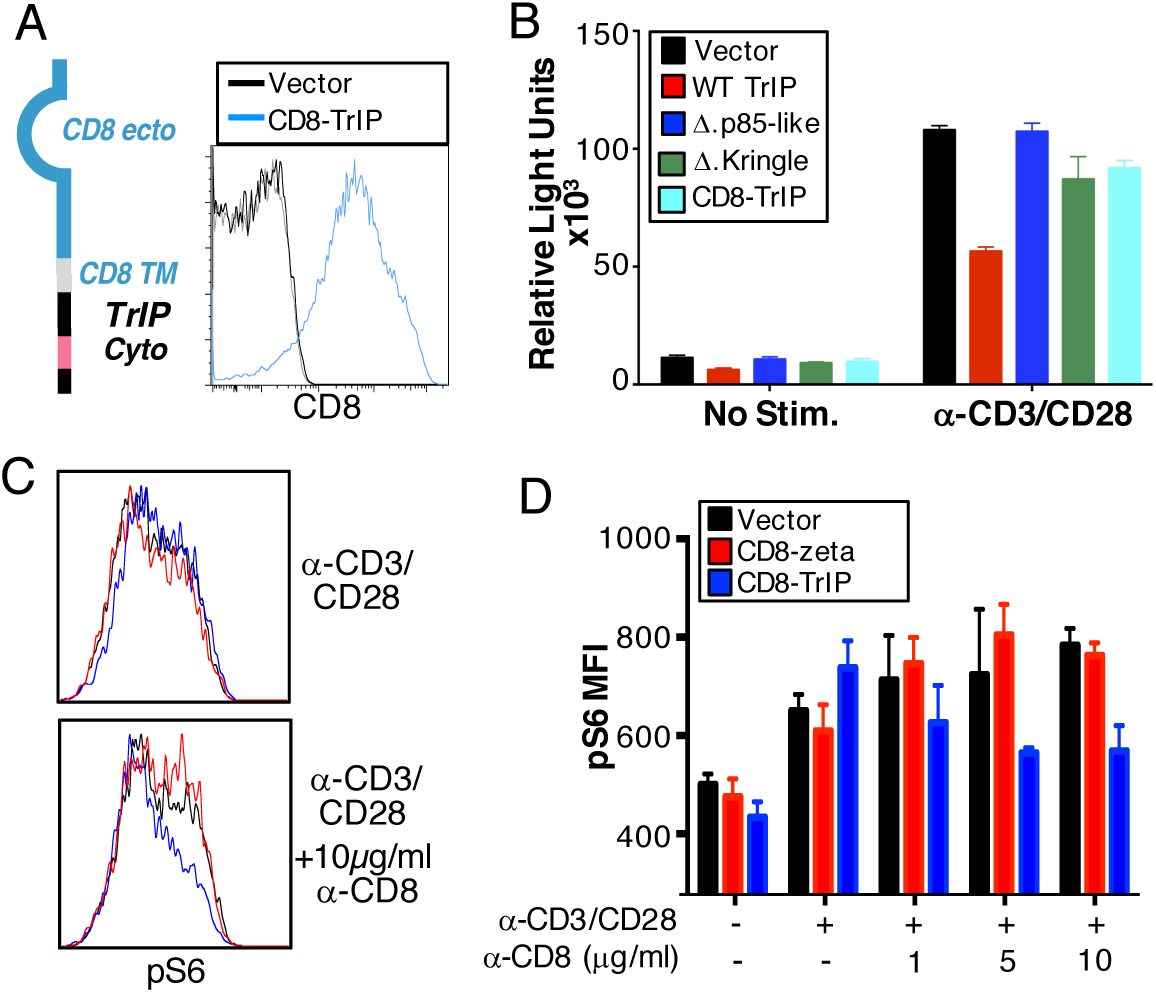
Artificial dimerization of TrIP induces inhibition of T cell activation. (**A**) Structure of ecto hCD8-mPIK3IP cytoplasmic chimera, and its expression on transfected D10 cells. (**B**) Luciferase assay on D10 cells transiently transfected with Flag-tagged TrIP, hCD8-TrIP and luciferase constructs and activated with anti-CD3/CD28. (**C**) Flow cytometric analysis of pS6 after stimulation of hCD8-TrIP transiently transfected D10 cells in the presence of 10 µg/ml anti-hCD8. (**D**) Quantification of pS6 after stimulation of hCD8-TrIP transfected D10 cells in the presence of varying concentrations of anti-hCD8. Data are representative of three experiments.

To determine whether inducible downregulation of TrIP only occurs in the context of stimulation by antigen and APCs (which may express a ligand for TrIP), we stimulated transfected D10 cells with α-CD3/CD28 and evaluated the surface expression of TrIP at various time points in the absence of APC and cognate antigen using the α-Flag antibody. Thus, we observed that even after CD3/CD28 Ab stimulation, without APC’s, WT TrIP expression was still downregulated (**Fig. 5A-B**) with a concomitant upregulation of pS6 (**Fig. 5B**). In contrast, neither Δcyto-TrIP (**Fig. 5A-B**), Δkringle-TrIP or Δp85-TrIP (**Fig. 5C**) expression was downregulated over the course of the stimulation by CD3 and CD28 Ab’s. These data suggest that one or more ligands responsible for TrIP inhibition of PI3K may be located on the T cells themselves and interacts with PIK3P1 in a kringle domain-dependent fashion. However, this does not rule out the presence of an identical or different ligand on the APC’s.

**Figure 5.**
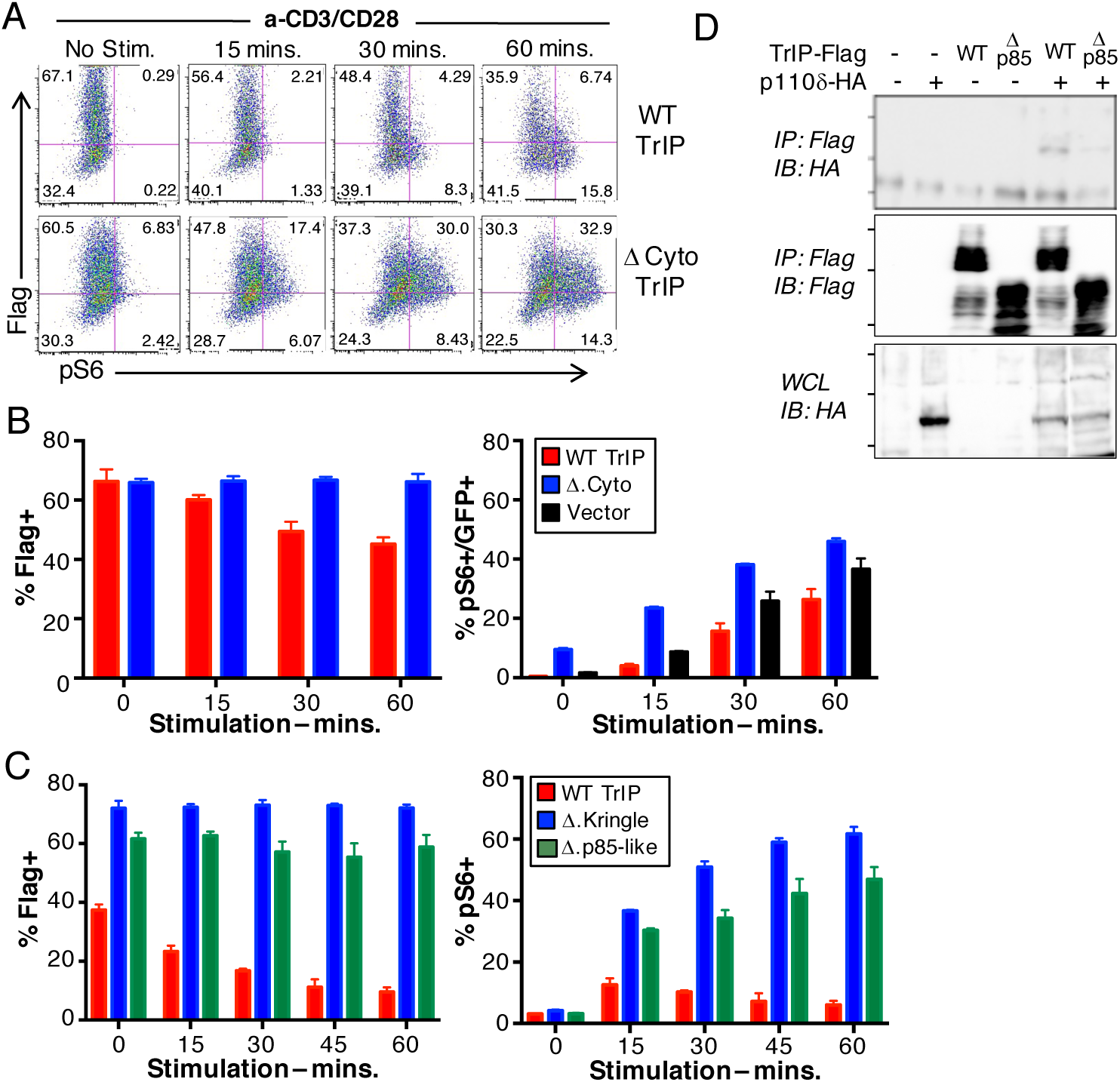
TCR signaling and TrIP function. D10 cells were transfected with control, WT TrIP or Δcyto-TrIP and stimulated with anti-CD3/CD28. (**A**) Representative flow cytometric analysis of Flag-tagged TrIP and pS6 expression of cells transfected with WT TrIP (top row) or Δcyto-TrIP (bottom row). (**B**) Quantitation of Flag expression over the course of stimulation; quantitation of total pS6^+^ cells at different time points. (**C**) TrIP surface expression (left) and pS6 staining (right) of cells expressing WT, Δ.kringle or Δ.p85-like TrIP, after stimulation with anti-CD3/CD28, as in panels A-B. (**D**) The cytoplasmic p85-like domain of TrIP binds to p110δ. 293 cells were transiently transfected with Flag-tagged WT TrIP and p85Δ-TrIP along with HA-tagged p110δ, as indicated. Co-immunoprecipitation and western blot analysis of TrIP and p110δ (top). Immunoprecipitated (IP) fractions showing Flag-tagged (TrIP) protein expression (middle). Whole cell lysate (WCL) showing HA-tagged (p110δ) protein expression (bottom). Data are representative of three experiments.

### The p85-like domain of TrIP interacts with p110δ PI3K

TrIP has been shown to interact with the PI3K catalytic subunits p110α/β in mouse embryonic fibroblast (MEF) cells, via a ‘p85-like’ domain with approximately 80% homology to the inter-SH2 domain of the PI3K regulatory subunit p85 (Zhu et al., 2007). However, it is not known whether TrIP can also interact with p110δ, which is the main catalytic subunit of PI3K activated by TCR signaling (Okkenhaug and Vanhaesebroeck, 2003). To test a possible interaction of p110δ with the p85-like domain of TrIP, we transiently transfected 293 cells with Flag-tagged WT TrIP or ∆p85-TrIP, along with HA-tagged p110δ, and performed co-IP and western blot analysis (**Fig. 5D, top panel**). Thus, WT TrIP could co-IP p110δ-PI3K; however, there was a significant reduction in the ability of ∆p85-TrIP to co-IP p110δ (**Fig. 5D top panel, last lane**). These results are consistent with a previous report that examined the interaction of TrIP with other p110 isoforms in fibroblasts (Zhu et al., 2007), and suggest that TrIP may inhibit T cell activation through effects on p110δ.

### Deletion of TrIP leads to dysregulation of primary T cell activation

In order to assess the function of TrIP in T cells *in vivo*, we generated mice with LoxP-flanked (floxed) *Pik3ip1* alleles (Pik3ip1^fl/fl^) and bred them to mice with a CD4-driven Cre recombinase transgene. Mice with CD4-Cre alone were used as controls. In the absence of a suitable TrIP antibody, T cells from spleen and lymph nodes of naïve homozygous (CD4-Cre x *Pik3ip1*^fl/fl^) and heterozygous (CD4-Cre x *Pik3ip1*^fl/wt^) TrIP conditional knockout mice were screened by RT-PCR for TrIP mRNA expression and compared to wild type mice (CD4-Cre *Pik3ip1*^wt/wt^) (**Fig. S1**). Analysis of thymus, spleen and lymph nodes from these mice showed similar percentages of CD4 and CD8 T cells in all compartments as well as normal numbers of nTregs (Foxp3^+^CD25^+^) suggesting that T cell development was normal (**Fig. S1**). Absolute cell numbers from these compartments were also similar across all strains (data not shown) as was the gross size of primary (thymus) and secondary (spleen and lymph nodes) lymphoid organs (data not shown). These data suggest that T cell-specific deletion of TrIP does not grossly affect T cell development or homeostasis.

Based on data shown above, we predicted that deleting TrIP in primary T cells would lead to enhanced TCR signaling. We thus stimulated lymphocytes from wild type (WT) and conditionally-deleted TrIP (KO) age-matched mice with α-CD3/CD28 for varying times and evaluated early T cell activation by pS6 and CD69 expression. Following stimulation, TrIP KO CD4^+^ and CD8^+^ T cells showed significantly higher pS6 at later time points (1-4 hrs.) (**Fig. 6A-D**). CD69, an early activation marker that can typically be observed approximately four hours after stimulation, was expressed more highly in KO T cells compared to wild type cells (not shown). These data are consistent with our observation of more robust phosphorylation of Akt at both T308 and S473 in peripheral T cells lacking TrIP (**Fig. 6E-F**). Taken together, these results demonstrate that the loss of TrIP in T cells promotes more efficient TCR signaling and early T cell activation, consistent with the effects of an inhibitory protein.

**Figure 6.**
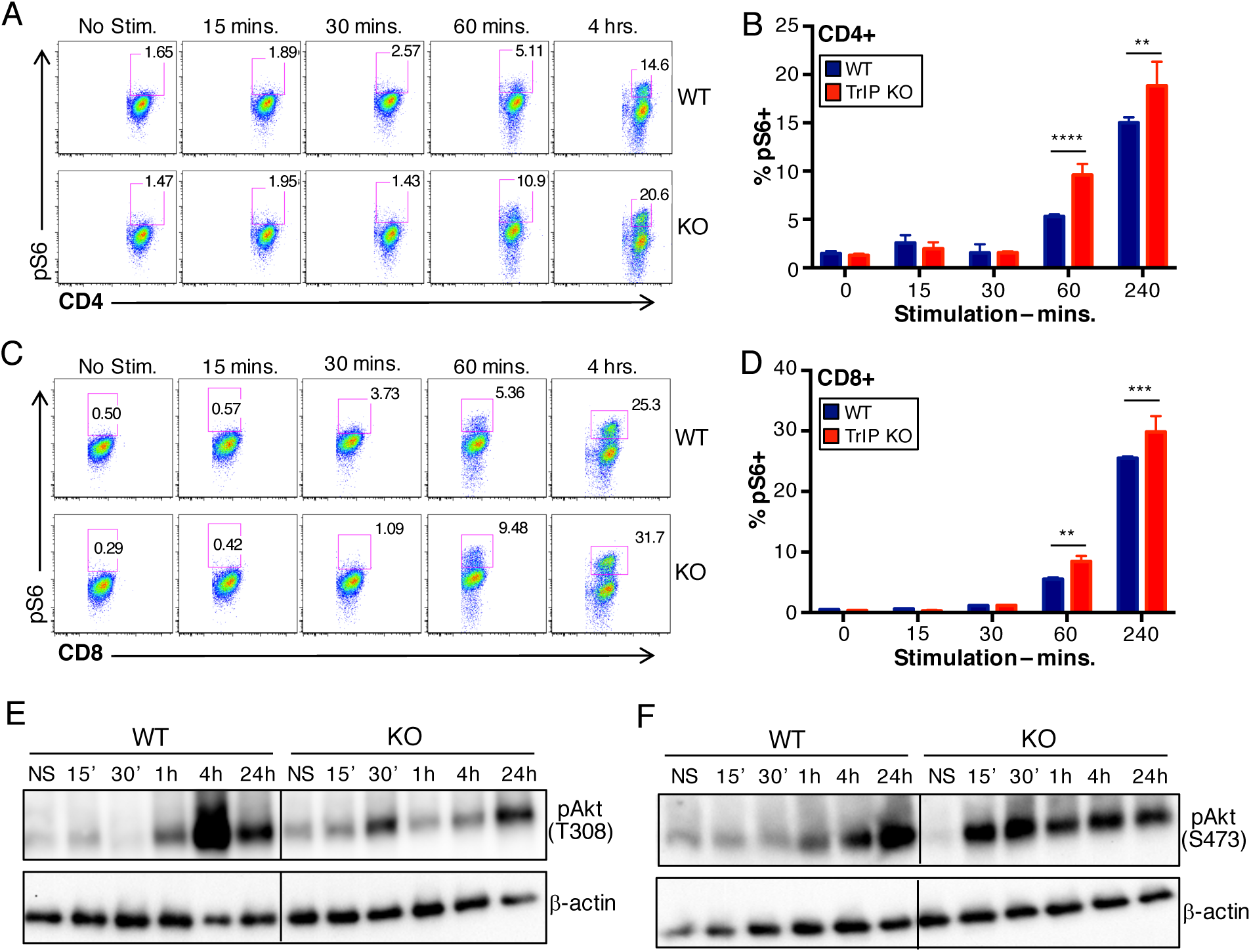
Conditional TrIP KO mice exhibit higher TCR signaling. Splenocytes and lymphocytes were isolated from WT and KO mice and stimulated with anti-CD3/CD28 for the indicated times. (**A**) Flow cytometric analysis of pS6 expression on stimulated CD4^+^ cells. (**B**) Quantitation of data shown in panel A. (**C**) Flow cytometric analysis of pS6 expression on stimulated CD8^+^ cells. (**D**) Quantitation of data shown in panel C. (**E-F**) T cells were purified from spleen and lymph nodes of WT or TrIP KO mice and stimulated with anti-CD3/CD28 antibodies for the indicated times. Lysates were analyzed by western blot for phospho-Akt (T308-left; S473-right) and β-actin as a loading control. *p* values were calculated using two-way ANOVA with Sidak’s multiple comparisons test. *p* values are represented with the following symbols: *0.01-0.05, **0.001-0.01, ***<0.001. Data in each panel are representative of at least three experiments.

### Deletion of TrIP enhances Th1 T cell polarization and inhibits iTreg generation

Previous research suggested that higher PI3K signaling may lead to enhanced Th1 potential, while lower levels of PI3K would favor generation of iTregs (He et al., 2008; Sauer et al., 2008). Based on our findings, we reasoned that T cell polarization towards a pro-inflammatory phenotype would be enhanced in TrIP KO T cells. We isolated naïve CD4^+^ T cells from WT and KO mice and cultured them with α-CD3/CD28 mAbs for three days under neutral (no polarization), Th0 (suppression of all polarization), Th1, Th17 and iTreg skewing conditions. At the end of culture, cells were re-stimulated with PMA/ionomycin for four hours and then evaluated for Th1 (T-bet and IFN-γ), Th17 (RORδt and IL17a) and iTreg (CD25 and Foxp3) phenotype. Thus, we noted a significant increase in the number of cells producing IFN-γ in TrIP KO T cell cultures, compared to WT cells, especially under Th0 and Th1, and even iTreg, conditions (**Fig. 7A**). Interestingly, we observed a smaller, but reproducible, decrease in the number of TrIP KO T cells making IL-17a, specifically under Th17 polarization conditions (**Fig. 7B**). More strikingly, we found that TrIP knockout T cells cultured under iTreg-generating conditions were significantly <u>less</u> efficient at producing Foxp3^+^CD25^+^ iTreg (**Fig. 7C-D**).

**Figure 7.**
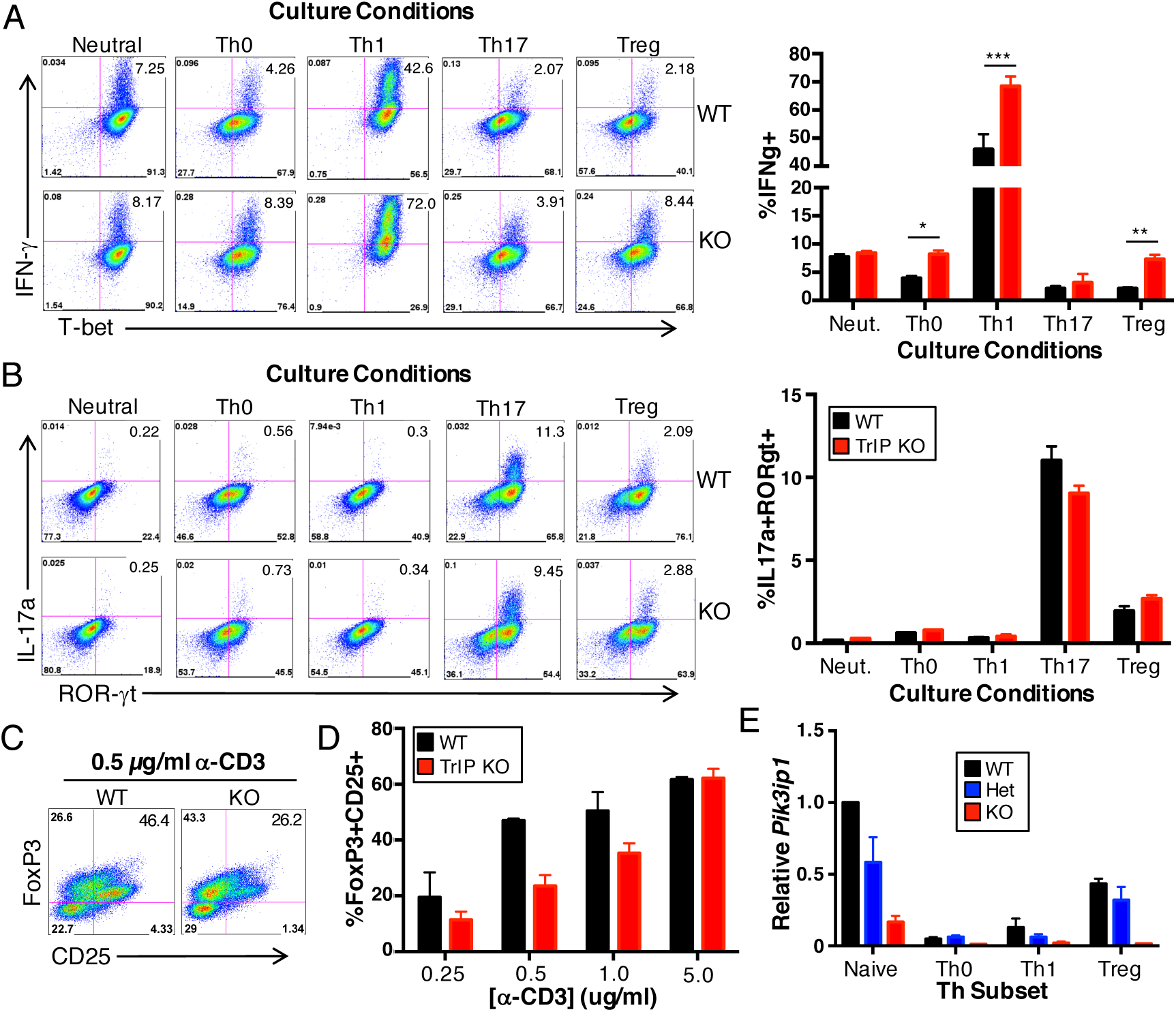
TrIP KO T cells display a higher tendency for Th1 differentiation. Naïve T cells from WT and KO mice were cultured under neutral, Th0, Th1, Th17 and iTreg conditions for three days, re-stimulated and analyzed for (**A**) IFNγ and T-bet expression, (**B**) IL-17a and RORγt expression, and (**C-D**) Foxp3 and CD25 expression. (E) Naïve CD4^+^ T cells from the indicated mice were stimulated under the indicated conditions, RNA extracted and analyzed by qPCR for *PIk3ip1* message. Data shown are normalized to naïve WT T cells. *p* values were calculated using two-way ANOVA with Sidak’s multiple comparisons test. *p* values are represented with the following symbols: *0.01-0.05, **0.001-0.01, ***<0.001. Data are representative of more than three experiments.

This result is consistent with previous findings that PI3K/Akt/mTOR signaling suppresses the generation of iTreg (Sauer et al., 2008). Interestingly, when we assessed the expression of *Pik3ip1* mRNA after culturing under the various Th polarization conditions, we found that *Pik3ip1* was severely downregulated in Th0 and Th1cells. By contrast, *Pik3ip1* was maintained at a somewhat higher level in iTreg, although still not as high as naïve cells (**Fig. 7E**), which is consistent with an apparent requirement to suppress PI3K signaling in Treg.

In order to determine whether the difference in Th1 polarization we had observed was actually due to the absence of PI3K modulation by TrIP, we performed the Th1 differentiation assay on WT and KO T cells with the addition of moderate concentrations of PI3K/Akt pathway inhibitors. Specifically, we employed an Akt1/2 inhibitor (Akt*i*-1/2), a pan-PI3K inhibitor (LY294002) and a selective p110δ inhibitor (IC-87114). Cells were stimulated with α-CD3/CD28 as in **Fig. 7** and cultured under Th1 conditions in the presence of varying concentrations of the PI3K inhibitors. At the end of culture, cells were re-stimulated with PMA/Ionomycin and evaluated by flow cytometry for IFN-γ expression. We found that IFN-γ production by KO cells was significantly inhibited at concentrations as low as 0.5 µM with all three of the inhibitors tested (**Fig. 8A-B**). Consistent with p110δ being the most prevalent catalytic subunit of PI3K linked to TCR signaling, IC-87114 was the most potent inhibitor at all concentrations. Overall, we observed a reduction of IFN-γ production by both WT and TrIP KO T cells in the presence of inhibitors, but this inhibition was more distinct with the KO cells. These results support the idea that the high IFN-γ production by TrIP KO T cells is due to a loss of TrIP inhibition of PI3K.

**Figure 8.**
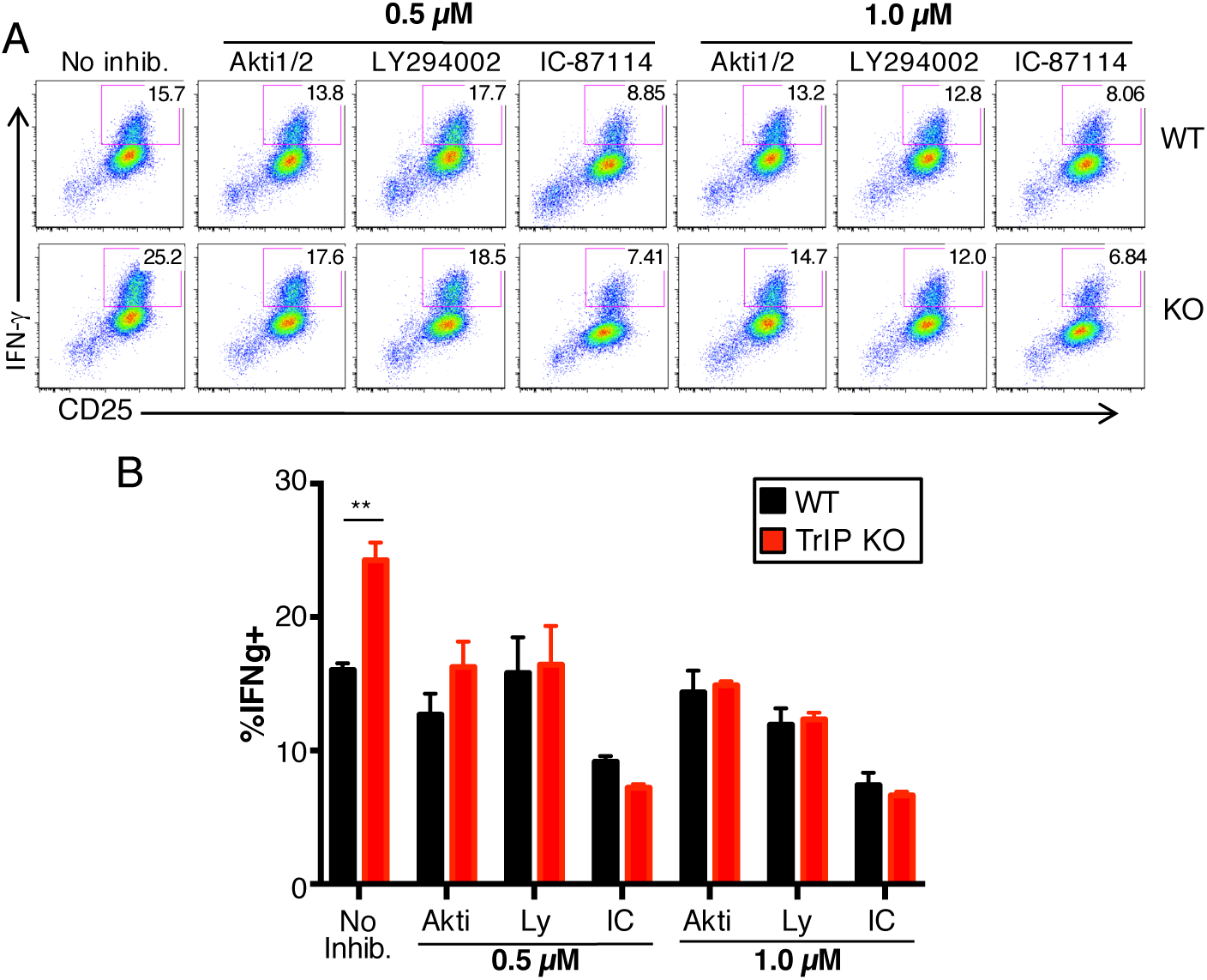
TrIP KO CD4^+^ T cell phenotype is reversed by PI3K/Akt pathway inhibitors. (**A**) Naïve T cells from WT and KO mice were cultured under Th1 conditions for three days in the presence of PI3K inhibitors (Akti, LY294002 and IC-87114), re-stimulated with PMA/Ionomycin and analyzed for IFNγ and CD25 expression. (**B**) Quantitation of A. *p* values were calculated using two-way ANOVA with Sidak’s multiple comparisons test. *p* values are represented with the following symbols: *0.01-0.05, **0.001-0.01, ***<0.001. Data are representative of more than three experiments.

### Enhanced in vivo function of T cells lacking TrIP

To explore the role of TrIP function in immune challenged mice, we infected mice with *Listeria monocytogenes*, a disease model that requires effective CD8^+^ and Th1 T cell responses for clearance and memory generation (Lara-Tejero and Pamer, 2004; Pamer, 2004). Specifically, we used a recombinant *Listeria monocytogenes* strain (LM-GP33) that expresses the GP(33-41) epitope from LCMV (Kaech and Ahmed, 2001). We infected mice with a relatively high titer (15,000 CFU) of LM-GP33, as we hypothesized that TrIP KO mice would be able to clear the infection more efficiently, based on the *in vitro* data detailed above. Thus, while all WT mice had culturable bacteria in the liver by day four after infection, all but one KO mouse had apparently cleared the infection by this point, at least below the limit of detection of the assay (**Fig. 9A**). The one KO mouse that did have detectable bacterial burden had significantly less than all other WT mice. We next evaluated the T cell response of these mice during the infection, and found that there was no significant difference in the percentage of LM-GP33-specific CD8^+^ T cells, based on tetramer staining (not shown). However, we did see a significantly higher percentage of short-lived effector cells (CD127^lo^KLRG1^hi^) and memory precursor effector cells (KLRG1^lo^CD127^hi^) within the CD8^+^ T cell compartment (**Fig. 9B**). Thus, the enhanced sensitivity of TrIP KO T cells described above does indeed translate to more robust *in vivo* activity of these cells in response to an intracellular bacterial infection.

**Figure 9.**
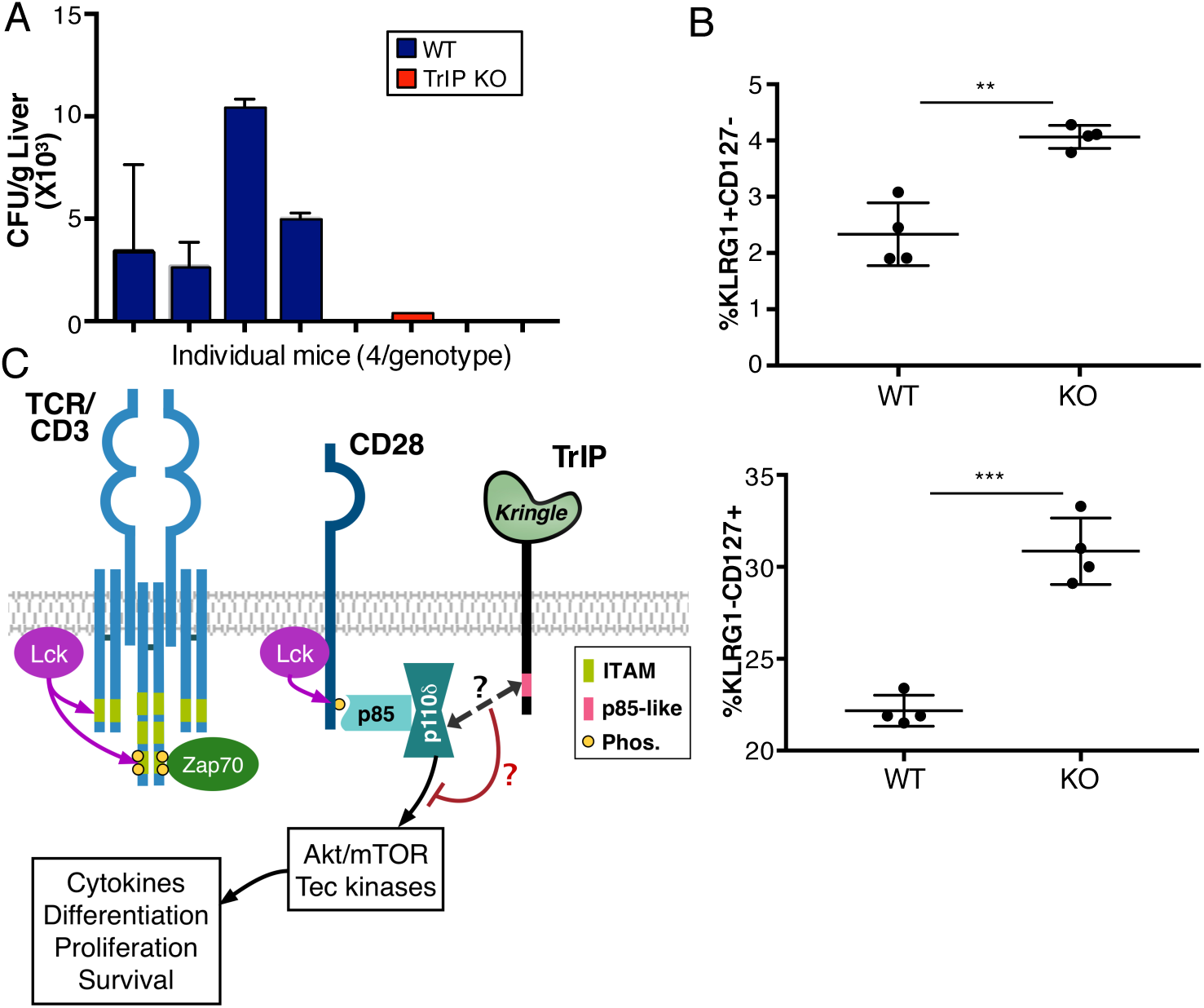
TrIP KO mice are less susceptible to Listeria infection. (**A**) Liver extracts from WT (CD4-Cre only) and TrIP KO mice infected with LM-GP33 were obtained four days after infection and plated to measure bacterial burden. (**B**) Analysis of tetramer positive CD8^+^ effector T cells four days after infection. Plots represent CD44^+^ KLRG1^+^CD127^−^ (top) and CD44^+^KLRG1^−^CD127^+^ (bottom) CD8^+^ T cells. (**C**) Model for the function of TrIP in T cells. Data are representative of two experiments with 4 mice per group.

## DISCUSSION

TrIP is a transmembrane protein that contains two identifiable domains: an extracellular kringle domain, implicated in ligand interaction in related proteins (Ji et al., 1998; Menhart et al., 1995; Patthy et al., 1984), and an intracellular ‘p85-like’ domain. An initial study showed that PI3K was inhibited by TrIP through its interaction with the catalytic subunit of PI3K via the p85-like domain (Zhu et al., 2007). We have also shown that silencing TrIP in T cells leads to an upregulation in T cell signaling (DeFrances et al., 2012). However, the role of the kringle domain in TrIP function remains largely unknown. With recent studies implicating a loss of function or expression of TrIP in some cancers (He et al., 2008; Wong et al., 2014), we sought to obtain a more complete understanding of TrIP structure and function in an immune context. In order to ascertain the molecular mechanism through which TrIP is regulated, we expressed Flag-tagged variants of TrIP in T cells. Our results showed that overexpression of WT TrIP in D10 cells resulted in a slower rate of phosphorylation of ribosomal S6 protein, further validating the observation that TrIP inhibits the PI3K/Akt pathway. We also obtained the first direct evidence that the p85-like domain is important for the inhibitory function of TrIP. Thus, in contrast to WT TrIP, ectopic expression of TrIP mutants lacking either the entire cytoplasmic region or just the p85-like domain did not inhibit T cell signaling.

In our studies, the inhibitory activity of TrIP was inversely correlated with surface expression of the protein, as only when TrIP expression was mostly downregulated, did we see a significant upregulation of pS6, suggesting TrIP expression must be downregulated in order for T cell signaling to proceed following TCR activation. While the mechanisms behind this downregulation are still not clear, there is precedent for acute down-regulation of other immune checkpoint molecules. Thus, Lag3 and Tim-3 can be inducibly cleaved from the cell surface by metalloproteases, resulting in enhanced T cell activation (Clayton et al., 2015; Li et al., 2007), while PTEN, an intracellular inhibitor of the PI3K pathway, is downregulated by multiple mechanisms, including post-translational modification and degradation (Hawse et al., 2015). At this point, we cannot rule out either one of these mechanisms in the downregulation of TrIP protein during T cell activation. In addition, we found that after long periods of stimulation, *Pik3ip1* message is also downregulated in effector T cells and, as we would expect, maintained at higher levels in iTreg.

It has been reported that kringle domains in other proteins can serve regulatory functions, including acting as sites for protein-protein interactions (Cao et al., 2002; Castellino and McCance, 1997; Thery and Stern, 1996; Tolbert et al., 2010). We therefore hypothesized that the kringle domain might serve to modulate the activity of TrIP. Indeed, as with the p85-like domain, the kringle domain appears to be essential for TrIP inhibitory function. Overexpression of a TrIP construct in which the kringle domain was deleted led to higher induction of pS6 (after either Ab or Ag/APC stimulation) than was observed in T cells transfected with WT TrIP or even cells transfected with empty vector, suggesting that this construct may function as a “dominant negative,” to inhibit endogenous TrIP. Furthermore, substitution of the kringle domain with the extracellular region of hCD8, and subsequent crosslinking by α-CD8, led to diminished induction of pS6 after TCR stimulation. We show that cell-surface TrIP expression is downregulated during the course of TCR stimulation, even in the absence of APCs, and only occurs with WT TrIP and not truncated variants, leading us to conclude that one or more ligands that activates TrIP inhibitory activity is present on T cells. How binding of a ligand, expressed by either an APC or the T cell itself, might regulate TrIP function is not clear at this point. One possibility is that the kringle domain is required for recruitment of TrIP into proximity of the TCR and/or CD28 during synapse formation. Another non-exclusive possibility is that ligand binding may cause a conformational change in TrIP which promotes the binding, and inactivation, of PI3K.

While we saw no obvious developmental defects in T cell compartments in mice with T cell-specific TrIP deficiency, we cannot rule out the possibility that TrIP may play a more subtle role in TCR repertoire selection. When stimulated, TrIP knockout T cells exhibited more robust, early activation and displayed higher Th1 and lower iTreg differentiation potential. The altered T cell lineage skewing in TrIP KO mice appeared to be a result (at least in part) of increased PI3K signaling, as PI3K inhibitors were able to reverse this phenotype. In addition, we hypothesized that the increased tendency of TrIP KO T cells to make IFN-γ could impact the immune response to an infection. Our analysis of LM-GP33 infected mice seven days after infection showed no significant differences between wild type and knockout mice (**data not shown**). This could be due to the fact that C57Bl/6 mice are generally able to clear *Listeria monocytogenes* and mount a peak T cell response by day seven. Our analysis at day four, however, showed that TrIP knockout mice had significantly higher percentages of both SLECs and MPECs, suggesting that in the absence of TrIP, these mice mounted a more robust T cell response. Previous work by others has shown that *Listeria* bacteria replicate for three days after infection of mice (Wong and Pamer, 2003). Here we observed that by day four, TrIP knockout mice had cleared the infection, compared to wild type mice that still had significant bacterial burden. Based on the typical immune response to *L. monocytogenes* infection (Pamer, 2004; Wong and Pamer, 2003), clearance of bacteria correlates with the peak of the immune response. It is possible that with the increased T cell signaling in the absence of TrIP, CD4^+^ T cells are able to recruit more APCs for priming of CD8^+^ T cells which results in faster clearance.

Overall, our results point to TrIP being a potentially attractive target for modulating immune responses under conditions of chronic infection, cancer or autoimmunity. In the former settings, briefly turning off or reducing the TrIP inhibitory pathway could allow for a more effective clearance of pathogens or tumors. Conversely, triggering an upregulation of TrIP activity or expression could help in combating pathological immune responses such as autoimmune diseases.

## ACKNOWLEDGEMENTS

We thank Sue Kaech (Yale), for providing LM-GP33 and Marie DeFrances (Dept. of Pathology) for helpful discussions and feedback on the manuscript. This work was supported by PHS grants AI126845 and AI095730 and by a supplement to grant PHS grant AI103022 (to LPK), and by startup funds from the University of Pittsburgh (to LDC). The authors have no conflicts of interest to disclose.

